# Evolving optimum camouflage with Generative Adversarial Networks

**DOI:** 10.1101/429092

**Authors:** Laszlo Talas, John G. Fennell, Karin Kjernsmo, Innes C. Cuthill, Nicholas E. Scott-Samuel, Roland J. Baddeley

## Abstract

We describe a novel method to exploit Generative Adversarial Networks to simulate an evolutionary arms race between the camouflage of a synthetic prey and its predator. Patterns evolved using our methods are shown to provide progressively more effective concealment and outperform two recognised camouflage techniques. The method will be invaluable, particularly for biologists, for rapidly developing and testing optimal camouflage or signalling patterns in multiple environments.

## Main

Historically, camouflage has been considered a prominent example of a prey versus predator arms-race^1^, whereby one species gradually evolves harder-to-see colouration which, as a consequence, exerts evolutionary pressure on the other species for a more effective detection system^2^. Despite the expectation that camouflage will become progressively more effective, it has been challenging to model how the evolution of optimal camouflage might take place in a particular environment^3^. This problem has inspired biologists for centuries, ever since Erasmus Darwin claimed that “the colours of many animals seem adapted to their purposes of concealing themselves either to avoid danger, or to spring upon their prey”^4^.

In recent years, research has focused predominantly on testing the advantage of particular camouflage strategies using predefined patterns designed by the experimenter^5^. Although these studies are able to provide strong evidence that certain camouflage works better than others, they have limited power to explain what would be the optimum pattern for concealment. One of the challenges is simply the number of potential patterns in a complex visual environment: the parameter space for all possible colour and texture combinations is often gigantic.

One solution to this problem is to employ dynamically evolving stimulus sets in detection experiments. Bond and Kamil presented blue jays with digital moths on computer screens in greyscale, with birds trained to peck on detected prey items^6^. The digital moths evolved on the basis of predetermined “genes”. While this approach was effective, improving survival, manually encoding genes for a specific task makes generalisability difficult: for example, increasing the parameter space beyond a certain complexity (using colour rather than greyscale, say) makes testing live subjects unrealistic because of the number of trials required. However, putting a credible *artificial* observer into the evolutionary loop would circumvent this problem.

Recently, methods that stem from Artificial Intelligence have proved capable of deceiving human observers: deep neural networks can mimic fine art^7^ or create photorealistic images based on text descriptions^8^. Here, for the first time, we report an unsupervised method to create biologically-relevant camouflaged stimuli based on Generative Adversarial Networks (GANs)^9^. GANs employ competing agents, usually modelled as deep neural networks, to perform a zero-sum game. In their original example, Goodfellow and colleagues illustrated the underlying idea of GANs using a competition between police and a counterfeiter. The objective of the police (discriminative network) was to distinguish between counterfeit and real money, whilst the counterfeiter (generative network) aimed to produce counterfeit money that the discriminative network would falsely identify as real. Both agents evolved over time: the police became more sensitive to fake money, while the counterfeiter produced more and more authentic-looking forgeries. As pointed out by Goodfellow et al, over time, and if such a pair of strategies exist, these two systems will become stable at a so-called Nash equilibrium: given the two agents, with complete knowledge of their opponent’s strategy, there is no possible improvement that can be made to their own. Nash Equilibria, form the basic building block of evolutionary game theory, the theory, proposed by Maynard Smith and Price^10^, where these Nash equilibria often correspond to evolutionary stable strategies. This arms race between a counterfeiter and the police mirrors antagonistic agents, like predator and prey, and is therefore of inherent biological interest.

In particular, predators evolve, or learn, to locate prey by detecting them against some background, while prey evolve to remain undetected using protective colouration. The objective of the predators is to distinguish visual input that contains prey from empty scenes. Meanwhile, the prey aims to achieve a visual signature that makes a scene containing them look empty to a predator. In this example, the discriminative network can be thought of as the visual system of the predator that evolves over time to more effectively detect prey, and the generative network represents the genotype of prey, where new generations can inherit properties of previous survivors and exhibit better camouflage.

To model the evolution of camouflage and produce increasingly difficult-to-see patterns, we implemented GANs to conceal triangular targets presented against images of ash tree (*Fraxinus excelsior*) bark, a complex texture (Fig. 1). Targets were extracted from each network after a set number of iterations and contrasted with two control patterns: the average colour of backgrounds, and a pattern developed through Fourier analysis (Fig. 2). Averaging the background is considered to offer “good” concealment^11^ and, as in our study, is often used in camouflage research as a baseline control^12^. The Fourier approach has previously been shown to be highly effective^13^, as has the related technique of log-Gabor wavelets when used to assess camouflage in targets such as ours^14^. To quantify difficulty, we measured the reaction time for human participants to detect the targets when displayed on a computer screen.

**Figure 1.**
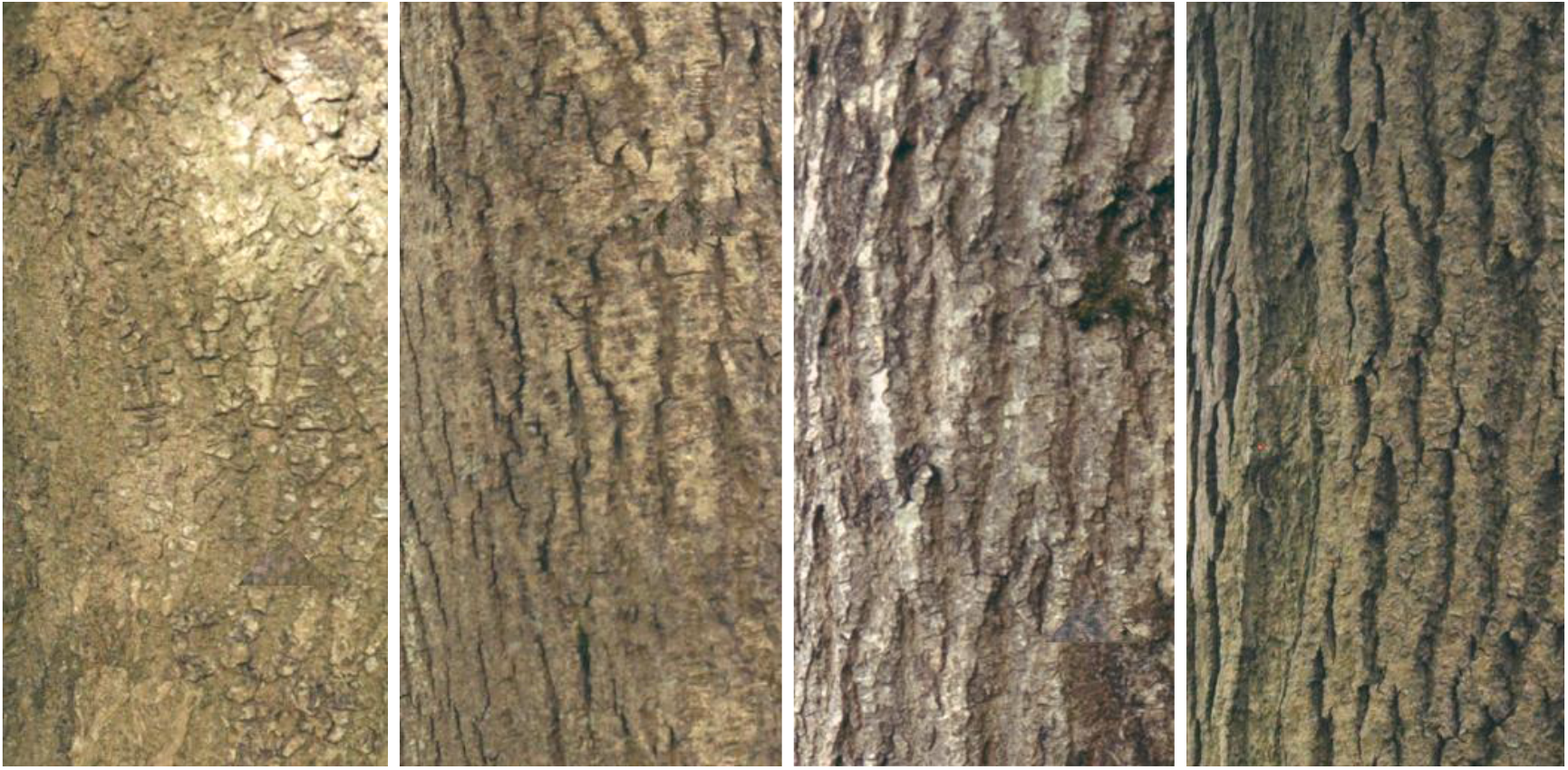
Examples of experimental stimuli. All examples feature targets evolved after 10,000 GAN iterations. See Figure S1 in the Electronic Supplementary Material (ESM) for revealed target locations.

**Figure 2.**
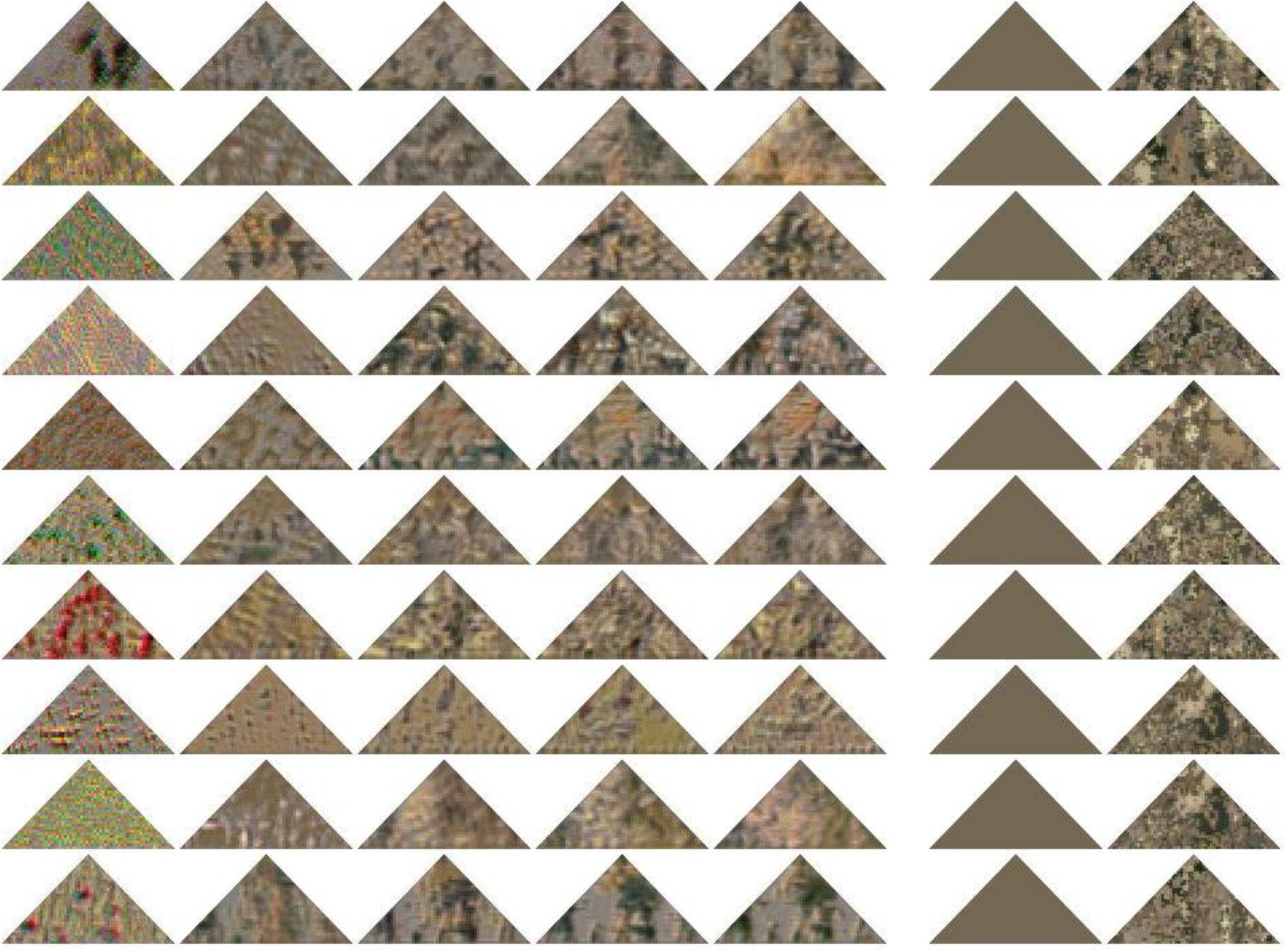
Example targets used in experiments. Columns 1-5 show targets evolved by GANs after 500, 2500, 5000, 7500 and 10000 iterations, respectively. Each row shows examples of targets from a different GAN. Columns 6 and 7 show examples of control targets: Average and Fourier, respectively.

It is important to note that contrary to other GAN implementations^15^, where the generative network modifies a whole image, in our implementation only the target was evolved by the generative network, leaving the background unmodified. Using this approach, we demonstrate that a purely artificial system can demonstrate the gradual evolution of camouflage.

We found that targets produced by GANs after more iterations were increasingly harder to find. In the first analysis, the effect of increasing training steps on reaction time was examined. General linear mixed models (GLMMs) were used to show that targets became significantly harder to find as the number of iterations increased. The effect of iterations on log-transformed reaction times were analysed by fitting general linear mixed models. Fitting the simplest model gave an estimate for the effect of training steps on reaction time of 2.077 × 10^-5^ (SEM = 1.006 × 10^-6^) and this was highly significantly different from zero (Δdeviance = 418.42, d.f. = 1, p < 0.0001). This result demonstrates that our method can successfully illustrate an evolutionary arms-race, producing camouflage that is increasingly difficult to identify. From visual inspection it is clear that the largest changes occur at earlier stages of pattern evolution with the rate of change in patterns beginning to decrease beyond 5,000 iterations (Fig. 2). Accordingly, increments in detection times also started to diminish (Fig. 3).

**Figure 3.**
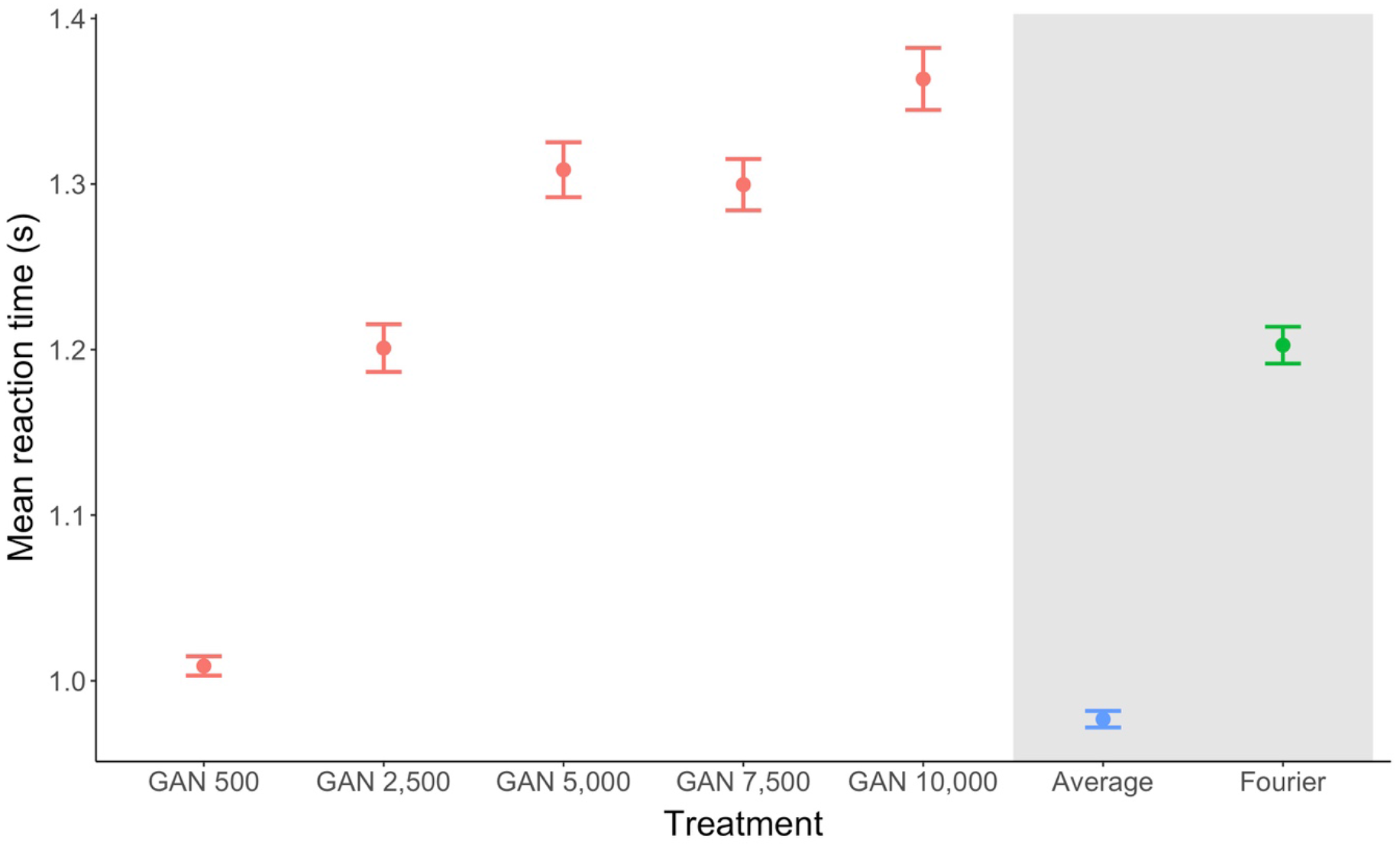
Mean reaction times for experimental stimuli. Error bars represent standard errors derived from a GLMM with participant as a random effect.

Furthermore, targets evolved by GANs were more effective than controls (Fig. 3). Treatment means were significantly different (Δdeviance = 1089.7, d.f. = 6, p < 0.0001). Based on Tukey post hoc tests all GAN-derived stimuli greater than 500 steps had significantly higher mean reaction times than Average targets. Fourier targets were significantly harder to detect than Average (p < 0.001) but GAN-derived stimuli with 5,000+ training step were significantly harder to detect than Fourier (p < 0.001). For details on the Tukey post hoc tests see Table S1 in the ESM.

We also found that some GANs produced more effective camouflage than others. Reaction times to GAN-derived stimuli of 10,000 training steps were selected and grouped by the network of origin. A random effects model with a common slope but different intercepts was chosen as the initial model. The effect of networks on reaction time was significantly different from zero (Δdeviance = 29.144, d.f. = 9, p < 0.0001). Mean reaction times ranged between 1.57 (SEM = 0.09) and 1.25 (SEM = 0.04) seconds (see Figure S2 in the ESM). In this study, both generator and discriminator networks were initialised with white noise, which is the reason why patterns at low iterations have a high inter-network variability (see first column in Fig. 2). We used this setup to demonstrate convergent evolution: the visual variance between the chosen backgrounds of tree bark was low and hence we expected that networks would come up with similar (and similarly effective) solutions after a higher number of training iterations. Nevertheless, certain networks were found to produce significantly harder to see patterns than others, which suggests that our method has the potential for modelling polymorphic scenarios, commonly found in nature^16^. The method can also clearly be adapted to use fixed initialisations, for example one could initialise the discriminator with pre-trained networks capable of better target detection^17^.

Our implementation follows a design that was deliberately simple, and we acknowledge that many alternative and more complex GAN architectures could be employed^18^. However, we believe that maintaining a simple architecture aids understanding and allows easier implementation for early adopters.

One promising development that could be beneficial in modelling biological systems is introducing multiple discriminator networks, standing for multiple observers influencing the target (generator network). For example, one of the discriminators could be limited to dichromatic representations of the target, simulating a typical mammalian predator^19^, or with altered visual acuity or viewing distance. It is also possible to introduce restrictions and limitations to the generator, other than the size and shape of the target; for example, bilateral symmetry.

We have demonstrated that GANs outperform other well-established methods for generating effective camouflage. This novel technique allows the exploration of high-dimensional feature and colour spaces in a way impossible using human, or non-human, observers. This obviously has applications for the development of military and civilian camouflage, but will also allow biologists to assess the trade-offs, beyond a pure concealment function, in natural camouflage patterns^20^. More widely, by reversing the reward function for the generative and/or discriminative networks, one can determine the optimal conspicuous signal and or sensory tuning for a given environment.

## Acknowledgements

LT and JGF were supported by an EPSRC grant (EP/M006905/1) awarded to NESS, RJB and ICC. We thank Jack Daniels and Thomas Ma for helping to take photographs of tree bark and grateful towards Erik Stuchly, Siyan Ye, Khishika Naidoo and Frankie King for their help during data collection.

## Author contributions

LT and JGF conceived and implemented the experiment, based on a framework devised by RJB, NESS and ICC. LT and KK collected the data. LT and JGF analysed the data. LT wrote the first draft of paper, with subsequent contributions by all authors.

These authors contributed equally: Laszlo Talas and John G. Fennell.

## Methods

### Participants

45 participants (4 male, 41 female) were recruited from the student population at the University of Bristol. The number 45 arises as a multiple of the number of generated ‘strains’ of GAN targets (see below). All participants had normal or corrected to normal vision. Informed consent was obtained from all participants as stated in the Declaration of Helsinki. All experiments were approved by the Ethics Committee of the University of Bristol’s Faculty of Science (application 60061) and were performed in accordance with relevant guidelines and regulations.

### Stimulus construction

We took 100 photographs of ash tree (*Fraxinus excelsior*) barks in October 2017 at Ashton Court Estate, Bristol, UK (2º64.8’ W, 51º44.6’ N). Images were taken from a distance of 1 m and a focal length of 18 mm using a Nikon D90 DSLR camera. Photographs contained an X-Rite ColorChecker Passport (X-Rite Inc., Grand Rapids, MI, USA), which was used to calibrate images using a cubic function implemented in Matlab 2016a (MathWorks 2016). Images were cropped so they only contained tree bark and resized to 1 pixel equalling 1.5 mm, using cubic interpolation.

Image size for the networks was selected to be 256 by 256 pixels, while the target triangle size was 32 by 64 pixels. Networks were trained on a custom-built PC with two graphical processing units (1x Nvidia Titan X and 1x Nvidia Geforce GTX 1080 Ti) using Keras (Chollet et al., 2015). The size of the training set was 3200 images, which comprised 32 randomly selected crops from each of the 100 images of ash bark.

The discriminative network was set to distinguish between empty scenes of tree bark and scenes with a target triangle present in the middle. To create effective camouflage, the task of the generative network was to modify the colour and texture of target triangles over randomly selected backgrounds so that the discriminative network would identify them as empty images.

Networks were trained for 10000 steps with a batch size of 32. The RMSprop optimiser was used for both the discriminative and generative networks, with learning rates of 2 × 10^-4^ and 1 × 10^-4^, and decays of 6 × 10^-8^ and 3 × 10^-8^, respectively. Binary cross-entropy was used as the loss function. The architecture of the discriminative network was: Conv2D(64), MaxPooling2d(2,2), Conv2D(128), MaxPooling2d(2,2), Conv2D(256), Conv2D(512), Flatten, Dense(1) and a sigmoid activation function to obtain predictions. All Conv2D layers had leakyReLU activations with alpha = 0.2 and ‘same’ padding. Dropouts were set to 0.5 for all Conv2D layers. The architecture of the generative network was: Architecture: Dense(8192, with dropout of 0.6), BatchNormalization, Dense(4096), BatchNormalization, Reshape(64,32,2), Conv2DTranspose(4,3), BatchNormalization, Conv2DTranspose(3,3) and a sigmoid activation function to normalise pixel values between 0 and 1. All batch normalisation had momentum of 0.9 and Conv2DTranspose layers had padding set to “same”. Ten networks were trained in total, with 15 evolved targets (strains) extracted after 500, 2500, 5000, 7500 and 10000 training steps from each network (Fig. 2), resulting in a total of 1050 GAN-derived targets.

In addition to the GAN targets, we included two control treatments: “Fourier” and “Average”. These were constructed using the following methods. Initially, 32 randomly positioned squares (sized 256 by 256 pixels) were cropped from each of the 100 images of tree bark. “Fourier” targets were constructed by decomposing the 3200 crops into energy and phase using 2-dimensional Fourier transformation, followed by taking pixel-wise average energy across the images. 15 targets were created by randomising the phase for each, and after an inverse Fourier transformation, the resulting images were indexed with 32 quantised colours obtained via minimum variance quantization and dithering of the original crops^13^. “Average” targets were created by taking the average colour of the same 3200 crops. Targets were created by cropping a 32 pixel high by 64 pixel wide triangle from the images. Both processes were repeated ten times and the resulting targets were grouped together with the GAN-derived targets, totalling seven treatment groups.

### Experimental procedure

A bespoke program, written using the Psychtoolbox-3 extensions^21,22,23^ for Matlab 2015b (The MathWorks, Inc., Natick, MA, USA) was used to construct and present the stimuli, and to collect experimental data. Each experimental trial consisted of a single target presented at a random position on randomly selected images of (ash) tree bark on a gamma-corrected computer display (Iiyama, Tokyo, Japan). The background images were 512 by 1024 pixels and subtended a visual angle of 26.5° by 53°. Targets had a size of 64 by 32 pixels, accounting for a visual angle of 5.5° by 2.75°. A central fixation cross on mid-grey background was displayed for 2 seconds prior to stimulus onset. To avoid spotting the target too early due a location close to the fixation cross, each target was placed at least 64 pixels away from the centre of the screen.

Participants were required to click on the detected target as quickly and accurately as possible, using a computer mouse. Their reaction times and whether they hit the target were recorded. Each trial had a 10 s time-out. Trials with timeouts and missed targets were removed from results.

Each participant was randomly assigned to a single strain of targets containing the seven treatment groups from all ten GANs, repeated five times in a random order, totalling 350 trials. Each of the 15 strains were exclusively presented to three participants only. In addition to the experimental trials, 10 practice trials using targets with a single random colour were presented to the participants at the beginning of the experiment to familiarise them with the task.

### Statistical analyses

GLMM analyses were initiated with the most complex model and were gradually simplified and assessed for significantly improved fits. Likelihood ratio tests were used to obtain p-values for the full model and the effect against a model without the effect. Nested models were compared using the change in deviance on removal of a term and by the Akaike Information Criterion. Analyses were carried out using the lme4 package^24^ in R^25^.

## References

1. Dawkins, R. & Krebs, J. R. Proc. R. R. Soc. B 205, 489–511 (1979).

2. Stankowich, T. & Coss, R. G. Proc. R. R. Soc. 274, 175–182 (2007).

3. Merilaita, S., Scott-Samuel, N.S. & Cuthill, I.C. Phil. Trans. R. Soc. Lond. B. 372, 20160341 (2017).

4. Darwin, E. Zoonomia; or, The Laws of Organic Life, London, Johnson (1794).

5. Troscianko, J., Skelhorn, J. & Stevens, M. BMC Evol. Biol.17, 7 (2017).

6. Bond, A. B. & Kamil, A. Nature 415, 609–613 (2002).

7. Gatys, L. A., Ecker, A. S. & Bethge, M. arXiv 1508.06576 [cs.CV] (2015).

8. Zhang, H. et al. arXiv 1612.03242 [cs.CV] (2017).

9. Goodfellow, I. J. et al. arXiv 1406.2661 [stat.ML] (2015).

10. Smith, J. M. & Price, G. R. Nature 246, 15–18 (1973).

11. Merilaita, S. & Stevens, M. Animal camouflage: mechanisms and function, Cambridge, Cambridge University Press (2011).

12. Cuthill, I. C. et al. Nature 434, 72–74 (2005).

13. Toet, A. & Hogervorst, M. A. Opt. Eng. 52, 041103 (2013).

14. Michalis, C., Scott-Samuel, N. E., Gibson, D. P. & Cuthill, I. C. Proc. R. Soc. Lond. B, 20170709 (2017).

15. Zhu, J-Y., Park, T., Isola, P. & Efros, A. A. arXiv 1703.10593 [cs.CV] (2017).

16. Karpestam, E., Merilaita, S. & Forsman, A. Sci. Rep.6, 22122 (2016).

17. Simonyan, K. & Zisserman, A. arXiv 1409.1556 [cs.CV] (2015).

18. Creswell, A. et al. IEEE Signal Process. Mag. 35, 53–65 (2018).

19. Jacobs, G. H. Phil. Trans. R. Soc. B 364, 2957–2967 (2009).

20. Barnett, J. B., Michalis, C., Scott-Samuel, N. E., & Cuthill, I. C. PNAS, 201800826 (2018).

## References for Methods

21. Brainard, D. H. Spat. Vis. 10, 433–436 (1997).

22. Pelli, D. G. Spat. Vis. 10, 437–442 (1997).

23. Kleiner, M., Brainard, D., Pelli, D. Perception 36(2007).

24. Bates, D., Machler, M., Bolker, B. & Walker, S. J. Stat. Softw. 67, 1–48 (2015).

25. R Core Team. R: A language and environment for statistical computing. R Foundation for Statistical Computing, Vienna, Austria (2017).

