## Supplementary Material for "Evolving optimum camouflage with Generative Adversarial Networks"

### Electronic Supplementary Material

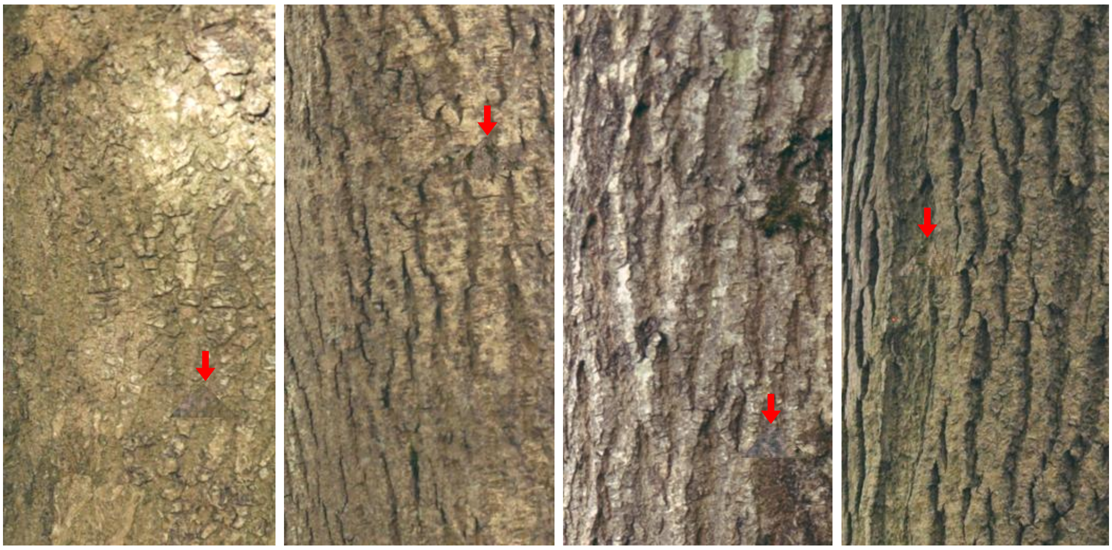

**Figure S1.** Revealed target locations of Figure 1 in the main article. A red arrow shows the locations of the targets.

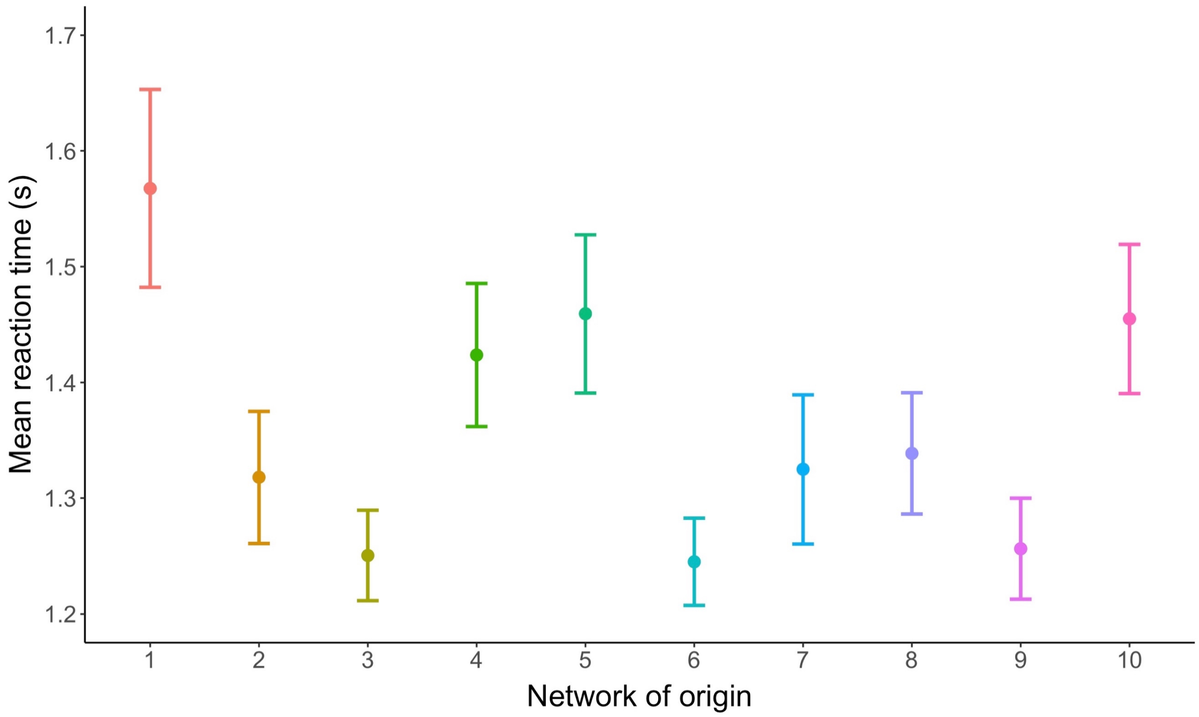


**Figure S2.** Mean reaction times for experimental stimuli from various Generative Adversarial Networks of origin. Error bars represent standard errors derived from a GLMM with participant as a random effect.

**Table S1.** Post hoc analysis of a General Linear Mixed Model, comparing all treatments pair-wise using Tukey HSD.

| Comparison | Estimate | Standard Error | z value | p |
|---|---|---|---|---|
| 2500 - 500 == 0 | 0.127 | 0.010 | 12.59 | < .001 |
| 5000 - 500 == 0 | 0.195 | 0.010 | 19.327 | < .001 |
| 7500 - 500 == 0 | 0.192 | 0.010 | 19.027 | < .001 |
| 10000 - 500 == 0 | 0.221 | 0.010 | 21.871 | < .001 |
| Fourier - 500 == 0 | 0.143 | 0.010 | 14.188 | < .001 |
| Average - 500 == 0 | -0.027 | 0.010 | -2.634 | 0.116 |
| 5000 - 2500 == 0 | 0.068 | 0.010 | 6.735 | < .001 |
| 7500 - 2500 == 0 | 0.065 | 0.010 | 6.439 | < .001 |
| 10000 - 2500 == 0 | 0.094 | 0.010 | 9.281 | < .001 |
| Fourier - 2500 == 0 | 0.016 | 0.010 | 1.599 | 0.683 |
| Average - 2500 == 0 | -0.154 | 0.010 | -15.215 | < .001 |
| 7500 - 5000 == 0 | -0.003 | 0.010 | -0.295 | 0.999 |
| 10000 - 5000 == 0 | 0.026 | 0.010 | 2.547 | 0.143 |
| Fourier - 5000 == 0 | -0.052 | 0.010 | -5.136 | < .001 |
| Average - 5000 == 0 | -0.222 | 0.010 | -21.948 | < .001 |
| 10000 - 7500 == 0 | 0.029 | 0.010 | 2.841 | 0.068 |
| Fourier - 7500 == 0 | -0.049 | 0.010 | -4.84 | < .001 |
| Average - 7500 == 0 | -0.219 | 0.010 | -21.648 | < .001 |
| Fourier - 10000 == 0 | -0.078 | 0.010 | -7.681 | < .001 |
| Average - 10000 == 0 | -0.248 | 0.010 | -24.49 | < .001 |
| Average - Fourier == 0 | -0.170 | 0.010 | -16.812 | < .001 |
